# Integrating multi-omics data through deep learning for accurate cancer prognosis prediction

**DOI:** 10.1101/807214

**Authors:** Hua Chai, Xiang Zhou, Zhongyue Zhang, Jiahua Rao, Huiying Zhao, Yuedong Yang

## Abstract

**Background:** Genomic information is nowadays widely used for precise cancer treatments. Since the individual type of omics data only represents a single view that suffers from data noise and bias, multiple types of omics data are required for accurate cancer prognosis prediction. However, it is challenging to effectively integrate multi-omics data due to the large number of redundant variables but relatively small sample size. With the recent progress in deep learning techniques, Autoencoder was used to integrate multi-omics data for extracting representative features. Nevertheless, the generated model is fragile from data noises. Additionally, previous studies usually focused on individual cancer types without making comprehensive tests on pan-cancer. Here, we employed the denoising Autoencoder to get a robust representation of the multi-omics data, and then used the learned representative features to estimate patients’ risks.

**Results:** By applying to 15 cancers from The Cancer Genome Atlas (TCGA), our method was shown to improve the C-index values over previous methods by 6.5% on average. Considering the difficulty to obtain multi-omics data in practice, we further used only mRNA data to fit the estimated risks by training XGboost models, and found the models could achieve an average C-index value of 0.627. As a case study, the breast cancer prognosis prediction model was independently tested on three datasets from the Gene Expression Omnibus (GEO), and shown able to significantly separate high-risk patients from low-risk ones (C-index>0.6, p-values<0.05). Based on the risk subgroups divided by our method, we identified nine prognostic markers highly associated with breast cancer, among which seven genes have been proved by literature review.

**Conclusion:** Our comprehensive tests indicated that we have constructed an accurate and robust framework to integrate multi-omics data for cancer prognosis prediction. Moreover, it is an effective way to discover cancer prognosis-related genes.

## Introduction

Cancer is a complex disease that involves a series of interactions between genes and environments. Patients of the same cancer type have been observed significantly variant in cancer outcomes among clinical studies, which contributes the most to hindering the development of effective therapies for cancers [1]. Therefore, it is important to precisely separate high-risk patients from low-risk ones according to genomic information. Currently, many studies have been designed to evaluate cancer prognosis risks based on genomics information [2], and the most frequently used data is gene expression (mRNA) measured by the microarray techniques [3]. With the development of next-generation sequencing techniques, many other types of genomic data are made available, including DNA methylation [4], miRNA [5], and copy number variation (CNV) [6]. Since these techniques provide different views of cancer patients, it is beneficial to integrate multi-omics data for capturing complexity in the cancer prognosis prediction.

Recently, The Cancer Genome Atlas (TCGA) organization has sequenced multiple types of omics data from more than ten thousand samples over 33 cancer types [7], enabling integrative cancer analyses based on multi-omics data. In this way, many statistical methods have been developed for different biological questions. For example, Rohart *et al*. designed a general package based on the sparse partial least square-discriminant analysis for omics data integration and extraction [8]; Mariette *et al*. used an unsupervised multiple kernel framework to predict breast cancer clinical outcomes [9]; Kim *et al*. designed a grammatical evolution neural network to evaluate ovarian cancer risks [10]; Ahmad proposed a hierarchical Bayesian graphical model that combined a Gaussian mixture model with an accelerated failure time model to detect clinically relevant biomarkers for breast cancer [11]; Corett combined the sparse correlation matrix estimator with the maximum likelihood estimator algorithm to identify differentially expressed genes [12]; Tong developed the HC-Cox method with a hierarchical clustering framework to extract multi-omics data for colon cancer prognosis prediction [13]. Though these studies have been carefully designed to integrate multiple omics data, the employed linear methods are limited to capture the representative features from thousands of variables. It is even worse if considering the high dimensions of heterogeneous variables during the processing of multi-omics data [14].

Recently, deep learning (DL) techniques have been shown with superior performance in dealing with nonlinear problems, and many DL-based methods were designed for cancer survival analysis. For example, Cheerla *et al*. developed DL-Cox method by inputting the multi-omics features into a deep neural network to estimate cancer outcomes [15]. Chaudhary *et al*. employed the Autoencoder to extract representative features and used the features for liver cancer subtype identification [16]. Following this study, Lee *et al*. used the Autoencoder to rebuild representative composite features from three types of omics data, and input the learned features into the Cox model to separate the high-risk patients of lung cancer (AE-Cox) [17]. Li *et al*. individually used Autoencoders for each omics features respectively, and finally combined the generated features to predict the prognosis of breast cancer (ContactAE-Cox) [18]. However, Autoencoder is fragile from data noises when learning representative features for input data [19]. Additionally, these previous studies usually focused on the individual cancer type without making comprehensive tests on pan-cancer.

In this study, we designed a new framework to integrate multi-omics data by the denoising Autoencoder for accurate cancer prognosis prediction (DCAP). By inputting the multi-omics data into the unsupervised denoising Autoencoder (DAE), we obtained the representative features for the high dimensional input data, and then utilized these learned features to accurately estimate cancer risks through the Cox proportional hazard model. This framework was comprehensively tested on 15 cancers from TCGA database. By comparison, DCAP averagely improved C-index values by 6.5% over previous methods. Considering the difficulty to obtain multi-omics data in practice, we further fitted the estimated risks by training XGboost models based on mRNA data only. The constructed XGboost models were shown to achieve an average C-index value of 0.627 with dozens of features. As a case study, the independent tests on three breast cancer datasets from the GEO indicated that the constructed model by XGboost can separate high-risk patients from low-risk ones significantly (C-index>0.6, p-values<0.05) by using mRNA only. Based on genes identified by the XGboost and differential expression analysis, we identified nine prognostic markers (*ADIPOQ*, *NPY1R*, *CCL19*, *MS4A1*, *CCR7*, *CALML5*, *AKR1B10*, *ULBP2*, and *BLK*) highly associated with breast cancer prognosis, among which seven genes have been proved by literature review.

## Materials and Methods

### Datasets

In this study, we downloaded cancer datasets from TCGA level 3 (https://tcga-data.nci.nih.gov/tcga/) through the R package “*TCGA-assembler 2*” v1.0.3[20] The datasets contained four types of multi-omics data: mRNA, miRNA, DNA methylation, and copy number variation (CNV), where “mRNA” was RNA sequencing data generated by the UNC Illumina HiSeq_RNASeq V2; “miRNA” was miRNA sequencing data obtained by the BCGSC Illumina HiSeq miRNASeq; DNA methylation data was generated by the USC HumanMethylation450, and CNV data was generated by the BROAD-MIT Genome wide SNP_6.

Since CNVs and DNA methylations reflect information on the sites representing millions of variables, we extracted their respective gene-level features by averaging the copy numbers of all CNV variations or the DNA methylations in CpG sites on each gene. For all four types of omics data, we processed the missing values following the previous study [16]. In each cancer data, we excluded features that were missing in more than 20% of the patients, and then excluded patient samples if they missed more than 20% of the remaining multi-omics features. Afterward, we excluded cancer datasets with fewer than 50 uncensored samples. For the left samples, the missing values were imputed based on the median values by using R package “*imputeMissings*” [21]. For the mRNA and miRNA data, the expression values were transformed through the *log* function. Afterward, all features were standardized to a mean of zero and standard deviation of one based on all cancer samples. Finally, we used the common features shared by all these 15 cancers that include 16160 mRNA features, 354 miRNA features, 20123 methylation features, and 23600 CNV features. It should be mentioned that our study used data only from cancer patients without involving data from any normal persons or patients of other diseases.

Three external breast cancer datasets were collected from the GEO database (https://www.ncbi.nlm.nih.gov) for independent tests. Among these, GSE2990 contains RNA-seq data and survival information of 126 breast cancer samples submitted by the Princess Margaret Cancer Centre. In GSE9195 we downloaded 77 breast cancer patients’ information, and GSE17705 contains 298 breast cancer patients’ data shared from Nuvera Biosciences. All the datasets were processed to remove the batch effects using the R package “*limma*” [22].

**Table 1.** The statistic information of used cancer data in TCGA

| Cancer | Uncensored | Total | Cancer | Uncensored | Total |
| --- | --- | --- | --- | --- | --- |
| BLCA (Bladder) | 151 | 336 | LUAD (Lung) | 155 | 441 |
| BRCA (Breast) | 78 | 613 | LUSC (Lung) | 135 | 329 |
| CESC (Cervical) | 67 | 290 | MESO (Mesothelioma) | 74 | 87 |
| COAD (Colon) | 59 | 255 | PAAD (Pancreatic) | 92 | 173 |
| ESCA (Esophageal) | 75 | 180 | SARC (Sarcoma) | 93 | 250 |
| HNSC (Head&Neck) | 168 | 404 | SKCM (Skin) | 212 | 447 |
| LGG (Brain) | 124 | 504 | STAD (Stomach) | 144 | 366 |
| LIHC (Liver) | 123 | 357 | Total | 1750 | 5032 |

### The architecture of DCAP method

As shown in Fig 1, the high-dimensional features of multi-omics data were input into a DAE network to obtain the representative features, which were then utilized to estimate patients’ risks through the Cox model. Considering the difficulty of obtaining multi-omics data in clinics, we further constructed the XGboost models by using mRNA data to fit the estimated risks. The constructed models were used to predict the cancer patients’ risks in the independent datasets. Besides, based on genes identified by the XGboost and differential expression analysis, we identified 9 prognostic markers highly associated with breast cancer prognosis.

**Fig 1.**
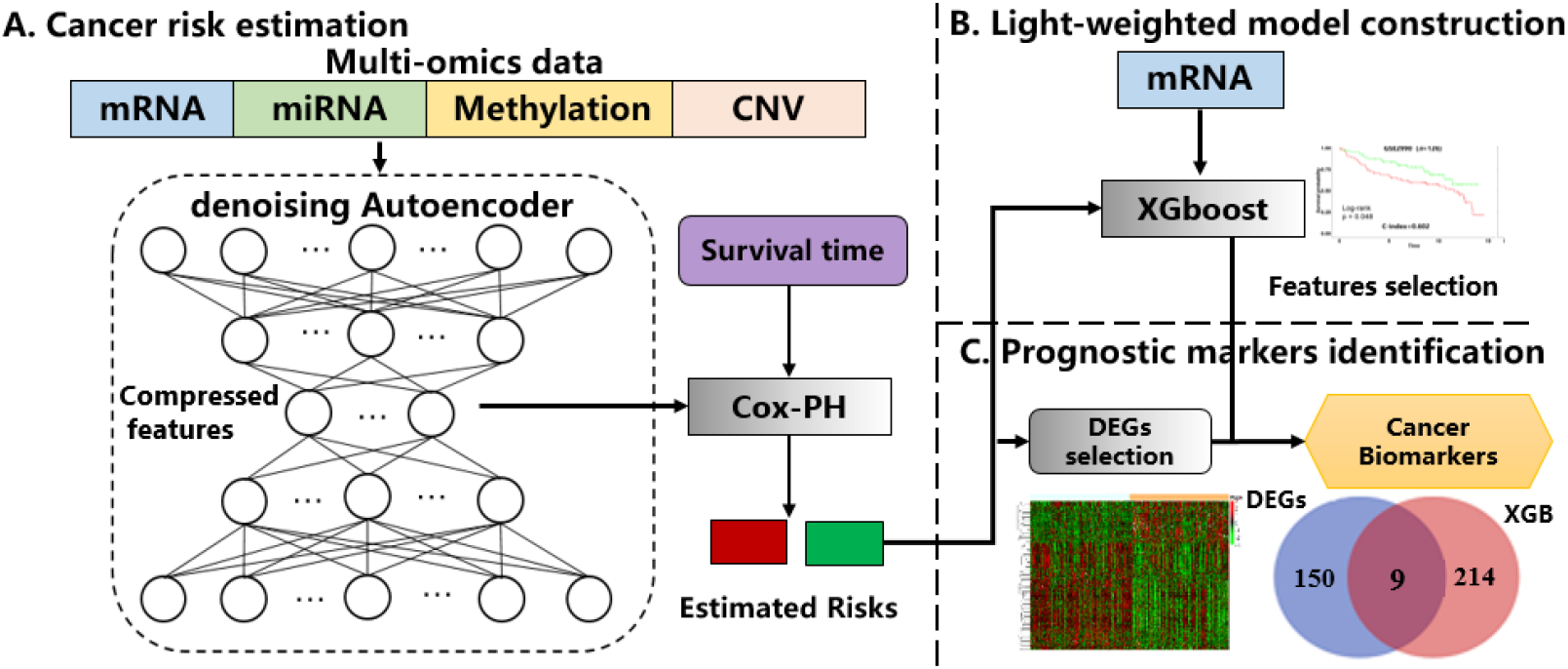
The architecture of DCAP proposed to integrate multi-omics data for cancer prognosis prediction. A) Estimating patients’ risks by using DCAP. B) Constructing XGboost models with mRNA data to fit the estimated risks. C) Identifying the prognostic markers highly associated with cancers.

### Denoising Autoencoder networks

Autoencoder is one kind of unsupervised neural network to learn an efficient representation of the input data. Supposing *x* = (*x*_1_,…, *x_n_*) is a list of input features, *x* is encoded to a smaller size of representative features that are decoded to *x*’, which is the output of the Autoencoder with the same size as *x*. The mean square error (MSE) was used to measure the difference between the input *x* and the output *x*’:

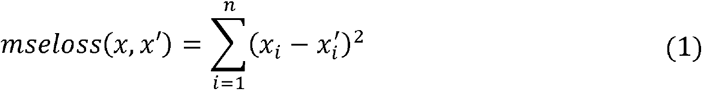

Compared with Autoencoder, DAE constructs the damaged data by adding noise to the high-dimensional features, and restores the original input by encoding and decoding steps. The design can make the deep neural network construct the real informative and robust low-dimensional representation. The damaged input is written as:

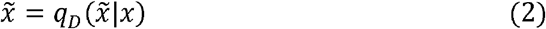

The loss of DAE is expressed as:

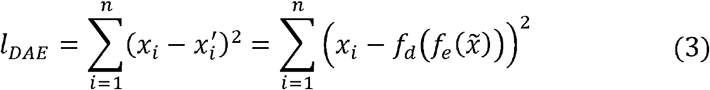

To avoid overfitting in our problem with high-dimensional features, we added an L2 regularization penalty term as

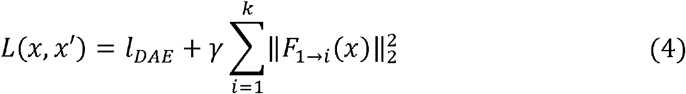

where *γ* is the coefficient for the L2-norm regularization penalty, *F*_1→*i*_ is the node activity in the deep neural network, *k* is the total number of layers (input, output, and hidden layers). Here, *γ* was set 0.0001, and *tanh* function was used as the activation function for all layers. In this study, considering the high-dimensional multi-omics features (>60000), we used a deep neural network with three hidden layers, i.e. *k = 5*. Based on the results of convergence analysis and parameter sensitivity study (Fig S1-S2), we set the three hidden layers as [500, 200, 500], and the training epoch as 100. The RMSE was converged after 50 epochs (Fig S1). The DAE was trained by back-propagation via the Adam optimizer. For different cancer types, we selected learning rate (LR) from {0.01, 0.001, 0.0001}, and the batch size from {32, 64, 128} based on the optimized loss values. Here, the decoded output 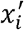 was used to guide the encoding of representative features, which will then be input into the Cox-PH model.

### Cox proportional hazard model for patients’ risk estimation

The learned representative features from the middle-hidden layer of the DAE network were used for building the Cox proportional hazard (Cox-PH) models to estimate the cancer risks. The multivariate Cox-PH model is defined as:

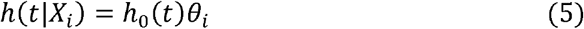

where *h*_0_(*t*) is the baseline hazard function to describe how the risk changed at time *t*, and *θ_i_* = exp (*βX_i_*) is used to describe how the hazard varied in response between the coefficient vector *β* and covariate vector *X*_i_ for patient *i*. The probability of the death for the patient *i* at the time *t_i_* is written as:

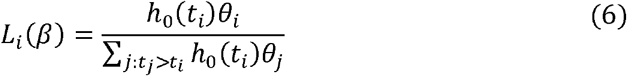

Hence, the corresponding *log* partial likelihood function is given as:

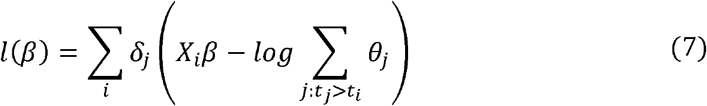

where *δ_j_* indicates the *j*th sample to be uncensored or not. This partial likelihood function was solved by using the Newton-Raphson algorithm as implemented by the “*glmnet*” package in R [23]. The computed *β* is used to estimate the risk scores in the Cox-PH model. At last, the patients are classified into two risk subgroups based on the median predicted risk value.

### mRNA-based XGboost model for risk prediction

To improve the interpretability and clinical applicability of our method, we employed XGboost to select a small number of key genomic features for building prediction models. XGboost is an ensemble of *K* regression trees (*T*_1_(*X*, *Y*)… *T_k_*(*X*, *Y*), where *X* is the feature vector and *Y* is the corresponding risk. Supposing that the dataset contains *n* examples and *p* features 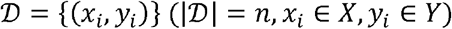, the ensemble XGboost model uses *K* trees to predict the patients’ risks:

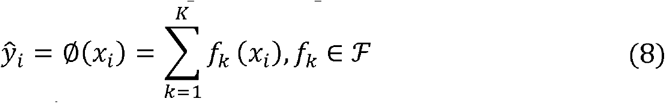

where 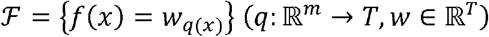 represents the space of the regression trees, *q* is the structure of the tree, *T* is the number of leaves in each tree, and *f_k_* represents a regression tree’s structure *q* with the weight *w*. This method was implemented by the “*XGboost*” package in R [24]. In this study, we selected the depth from [2, 8] and the learning rate from nine values (0.01 and 0.05*[1, 8]). These parameters were optimized by minimizing the mean square error through the 10-fold cross-validation (CV). All other parameters in our study used the default values in the R package “*XGboost*”. The selected genes by XGboost were considered candidate genes related to cancer patients’ survival.

### Evaluations of cancer prognosis prediction

In the cancer prognosis prediction, the performance was usually estimated through the C-index values. The C-index represents the fraction of all pairs of individuals whose predicted survival times are correctly ordered based on the Harrell’s C statistics [25]:

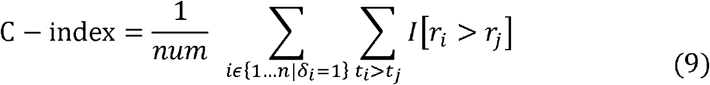

where *r_j_* and *r_j_* are the predicted survival risks for patients *i* and *j*, *t_i_* and *t_j_* are the actual survival times for patients *i* and *j*, *δ* indicates the sample is uncensored or not, *num* denotes the number of comparable patient pairs and *I*[.] is the indicator function. A higher C-index value indicates a better prediction with a value of 0.5 meaning a random prediction. In addition, the log-rank p-value was computed by the “*survival*” package in R for the probability to better separate patients into high-risk and low-risk groups by random.

### Differential expression analysis

Differential expression analysis is a kind of downstream analysis after cancer prognosis prediction, which is used for identifying genes with the most significant expression differences according to the divided sub-groups. These genes are regarded as potential targets that may affect the prognosis of cancer patients. In our study, the differentially expressed genes (DEGs) were genes with |log_2_(fold change)| > 0.5 and corrected p-value <0.05 detected by the “*limma*” package in R [22]. At last, the DEGs which were also selected by XGboost are seen as the prognostic markers which highly related to cancer prognosis.

### Parameters optimization and model training

For each cancer type, we randomly split the TCGA data into 80% for training and 20% for test. Each time, the representative features were reconstructed by the DAE network, and the hyperparameters were optimized according to the loss function. For the training of Cox and XGboost models, hyperparameters were optimized based on the 10-fold CV in the training set. With the optimized parameters, a model was re-trained on the whole training dataset and tested on the test dataset for the final performance. To remove fluctuations brought by random selections of the test dataset, we employed a bootstrapping strategy and repeated this process 10 times to obtain averages.

### Methods for Comparisons

In this study, we compared the cancer prognosis prediction performance with three traditional methods (the Cox model used the reconstructed features by PCA (PCA-Cox), Cox with elastic net (Cox_EN) [26], and HC-Cox) and three deep learning-based methods (DL-Cox, AE-Cox, and ContactAE-Cox). We used default parameters for these methods.

## Results

### Patients’ risks estimation by multi-omics data

As shown in Table 2, DCAP achieved essentially the same C-index values for the 10-fold CV and independent tests with average values of 0.678 and 0.665 over 15 cancers, respectively. The close results indicated the robustness of our method. For the 15 cancer types, the C-index values ranged from 0.591 to 0.823 with the highest value for LGG and the lowest one for STAD. The LGG has the highest C-index value likely because LGG has a large sample size (the 2nd largest in the dataset).

**Table 2.** The C-index of the cross-validations and tests on 15 cancers by DCAP

| Cancer | Validation | Test | Test (95% confidence) | Cancer | Validation | Test | Test (95% confidence) |
| --- | --- | --- | --- | --- | --- | --- | --- |
| BLCA | 0.678 | 0.646 | 0.611~0.68 | LUAD | 0.649 | 0.629 | 0.574~0.642 |
| BRCA | 0.684 | 0.662 | 0.615~0.707 | LUSC | 0.581 | 0.597 | 0.540~0.651 |
| CESC | 0.69 | 0.685 | 0.651~0.718 | MESO | 0.792 | 0.765 | 0.720~0.809 |
| COAD | 0.663 | 0.622 | 0.566~0.676 | PAAD | 0.654 | 0.665 | 0.625~0.704 |
| ESCA | 0.607 | 0.594 | 0.549~0.638 | SARC | 0.731 | 0.719 | 0.683~0.753 |
| HNSC | 0.631 | 0.628 | 0.582~0.671 | SKCM | 0.656 | 0.644 | 0.590~0.695 |
| LGG | **0.832** | **0.823** | **0.795~0.843** | STAD | 0.575 | 0.591 | 0.551~0.629 |
| LIHC | 0.733 | 0.710 | 0.655~0.763 | Average | 0.678 | 0.665 | 0.620~0.705 |

We further detailed the contribution of each omics type in DCAP. As shown in Table 3, when using a single type of omics data, the mRNA performed best with an average C-index value of 0.628, and CNV had the lowest performance with a C-index value of 0.570. The miRNA and methylation ranked the 2nd and 3rd, respectively. Consistently, when excluding one omics type from the DCAP, mRNA caused the largest decrease of C-index value from 0.665 to 0.631, and the smallest decrease was caused by the exclusion of CNV. These results indicated that mRNA played the most important role in discriminating high-risk patients while CNV made the least contribution. On average, the prognosis prediction using multi-omics improved the C-index value by 5.9% over the one using only mRNA data.

**Table 3.** The contribution of each omics data for cancer outcome evaluation by using individual type of omics data or excluding one type from the final model

| Omics Type<sup>a</sup> | C-index | Omics Type<sup>b</sup> | C-index |
| --- | --- | --- | --- |
|  |  | DCAP | **0.665** |
| mRNA | 0.628 | -mRNA | 0.631 |
| miRNA | 0.617 | - miRNA | 0.639 |
| Methylation | 0.604 | - Methylation | 0.647 |
| CNV | 0.570 | -CNV | 0.654 |
| <sup>a</sup> Performances by using each omics type; <sup>b</sup> Performances by removing each omics type from the final model. |  |  |  |

### Comparisons with other methods

We compared the cancer prognosis prediction performance obtained by our method (DCAP) with other common methods using multi-omics data. As shown in Table 4, DCAP achieved the highest C-index values between 0.591 (STAD) and 0.823 (LGG), with an average of 0.665. Compared with other methods, DCAP improved C-index values by 6.5% on average. For other methods, the PCA-Cox method achieved the lowest C-index values with an average of 0.584, the other two traditional methods Cox-EN and HC-Cox achieved average C-index values of 0.602 and 0.615, which are lower than the DL-based methods. ConcatAE performed better than AE-Cox, but worse than our method. We implemented ConcatAE-Cox by using four types of omics data and achieved a C-index value of 0.658, which was slightly higher than their own reported one (0.644) that used two omics data (the methylation and miRNA)[18]. We also conducted the t-test on the results obtained by DCAP and the other methods, and the p-values demonstrated that our method had significant improvements over the other methods.

**Table 4.**
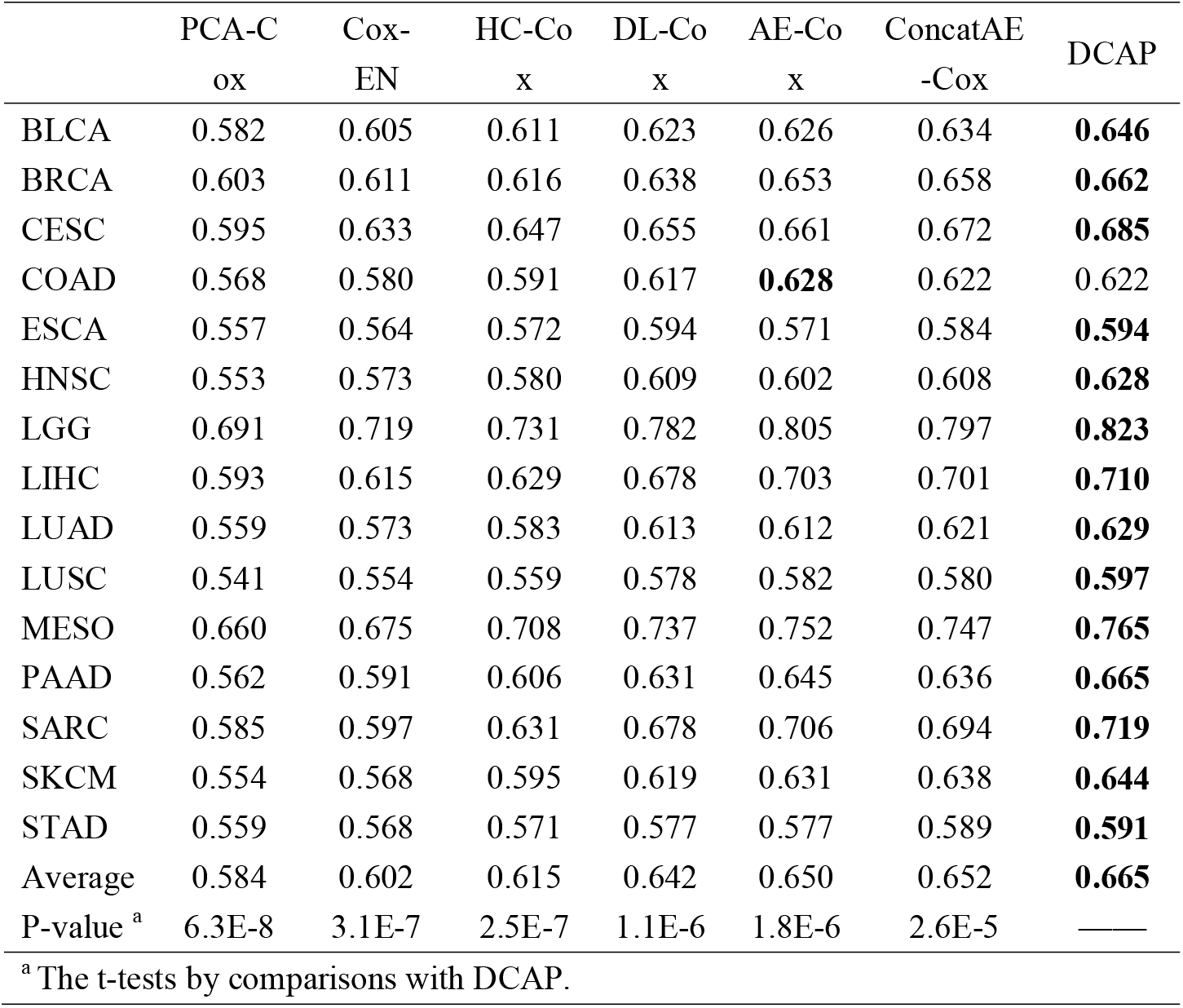
Methods comparisons by C-index values achieved on 15 TCGA cancers.

### Building light-weighted cancer risk prediction models by XGboost

The deep learning-based cancer prognosis prediction models constructed by DCAP are not interpretable without providing essential gene features. To extract important features, we employed the XGboost method to construct light-weighted models (DCAP-XGB). As shown in Fig 2, DCAP-XGB achieved C-index values between 0.565 (LUSC) and 0.755 (LGG), with an average of 0.627. The differences between the C-index values obtained by DCAP-XGB and DCAP ranged from 3.25% (BLCA) to 8.27% (LUAD), with an average of 5.64%. These results indicated that although feature selection caused a decrease of prediction, the XGboost can obtain comparable results with previous methods. More importantly, the XGboost model could make predictions with a small number of genomic features. As shown in Table S2, the models required 171-564 features with the most features for SKCM and the least ones for LUSC.

**Fig 2.**
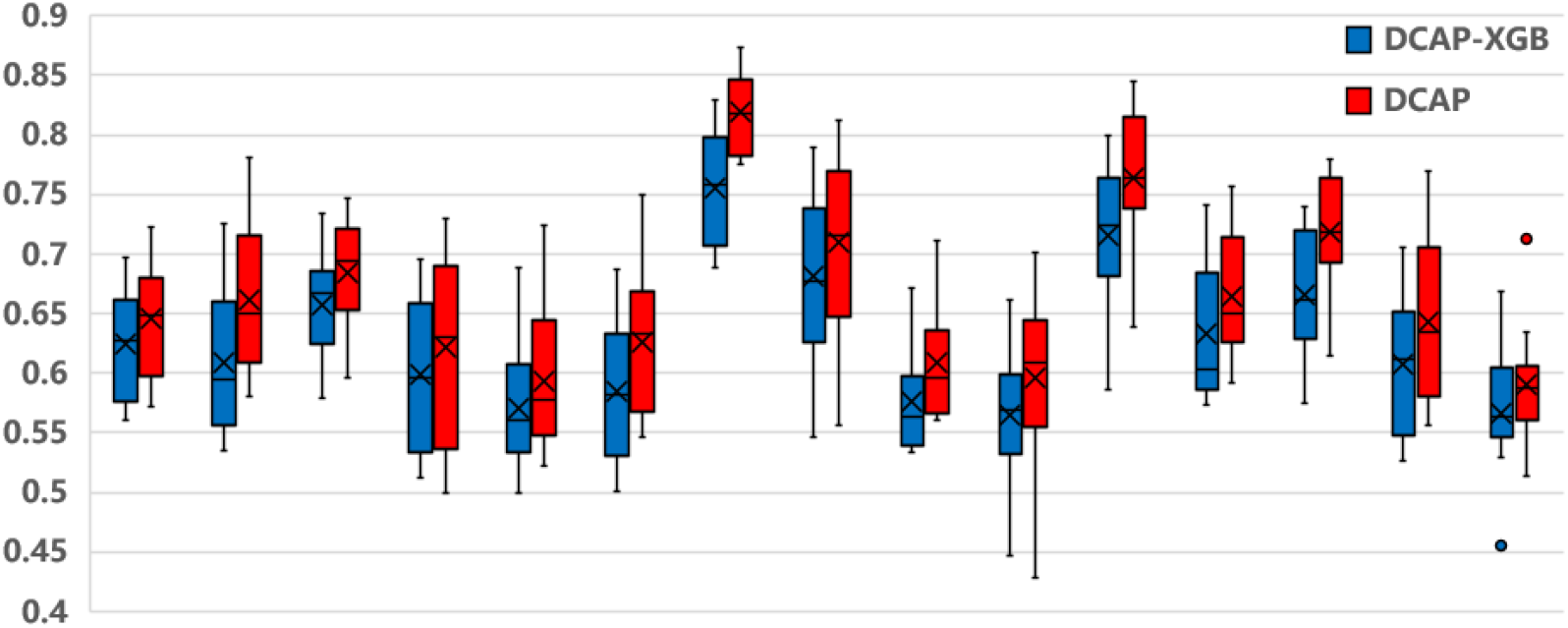
The C-index obtained by light-weighted DCAP-XGB compared with the DCAP. The blue parts are the C-index values obtained by DCAP-XGB and the red ones are those obtained by DCAP.

### Case study on the breast cancer

As a case study, we applied our method to the breast cancer (BRCA) that contains the largest number of samples. To validate the cancer prognosis prediction model constructed by DCAP-XGB, we tested the model on three external breast cancer datasets collected from the GEO database: GSE2990, GSE9195, and GSE17705. As shown in Fig 3A, for the three datasets, the predicted high and low-risk groups can be significantly separated from the survival curves with the p-values all below 0.05 and the similar C-index values (0.602, 0.605, and 0.611). These results indicated the robustness of our light-weighted risk prediction models.

**Fig 3.**
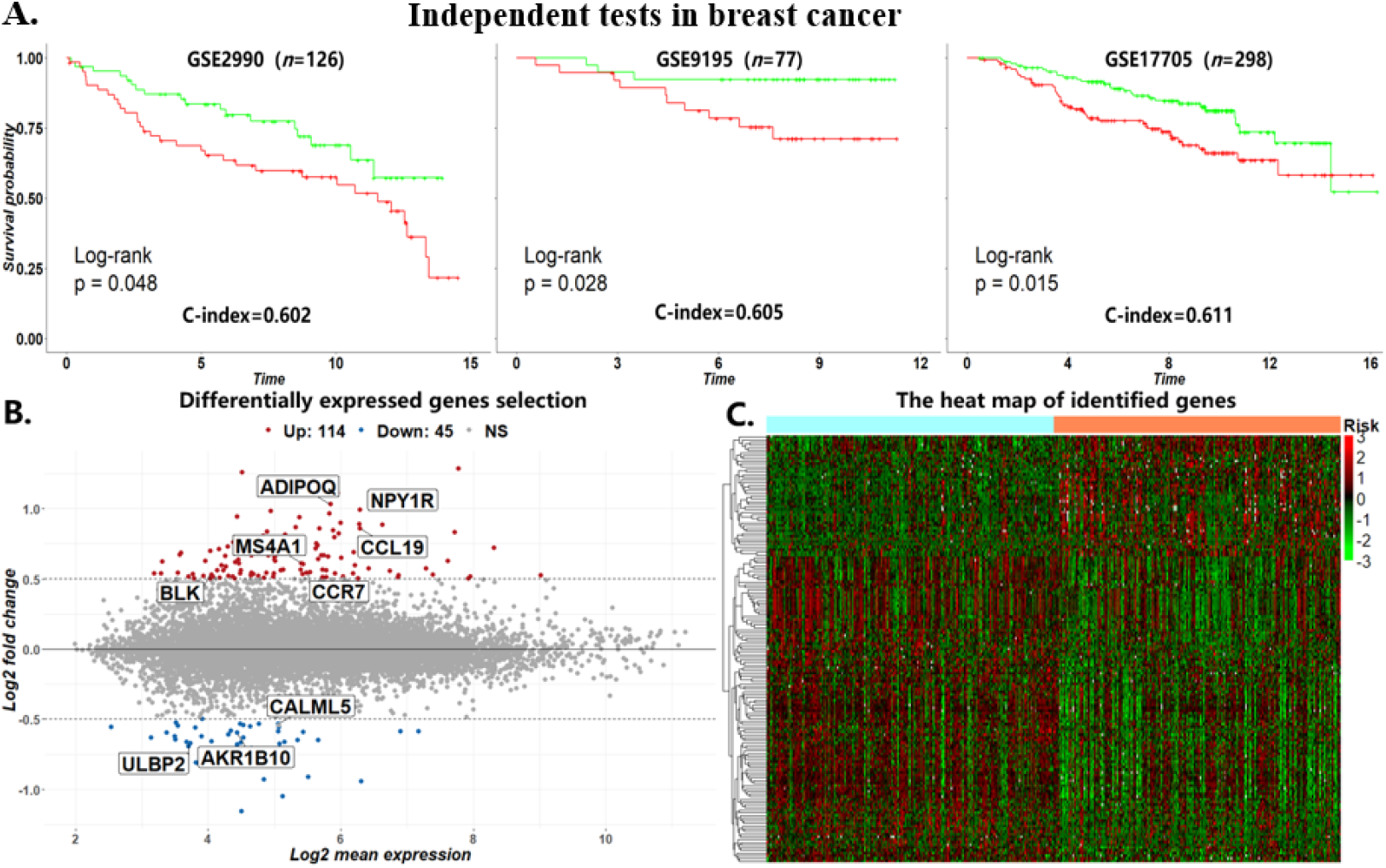
The case study on breast cancer. A) The results of the independent tests in three breast cancer datasets collected from GEO. B) The differentially expressed gene selection results in breast cancer. The red nodes represent the up-regulated risk genes and the blue nodes represent the down-regulated risk genes. The grey ones are the unselected genes. C) The heat map of identified DEGs in breast cancer (corrected p-value <0.05 and |log2FoldChange| >0.5).

Based on the divided high-risk and low-risk groups by DCAP, we identified 159 DEGs with corrected p-value<0.05 and |log2 fold change| >0.5, among which there were 45 down-regulated risk and 114 up-regulated risk genes (Fig 3B). Fig 3C shows the heat map on the expression of the DEGs. Among these 159 DEGs, 57 (35.9%) genes were confirmed to relate to the breast cancer by literature review. When mapped with 223 genes selected by the XGboost model, nine DEGs were overlapped, and seven (77.8%) of these nine genes (*ADIPOQ*, *NPY1R*, *CCL19*, *MS4A1*, *CCR7*, *CALML5*, and *AKR1B10*) have been indicated to associate with the breast cancer (Table 5). For the remained two genes (*ULBP2* and *BLK*), although no literature has directly demonstrated an association with the prognosis of the breast cancer, the induction of *ULBP2* was reported to associate with pharmacological activation of p53 triggers anticancer innate immune response [27], and *BLK* is a true proto-oncogene capable of inducing tumors, which is suitable for studies of BLK-driven lymphomagenesis and screening of novel *BLK* inhibitors in vivo [28].

**Table 5.**
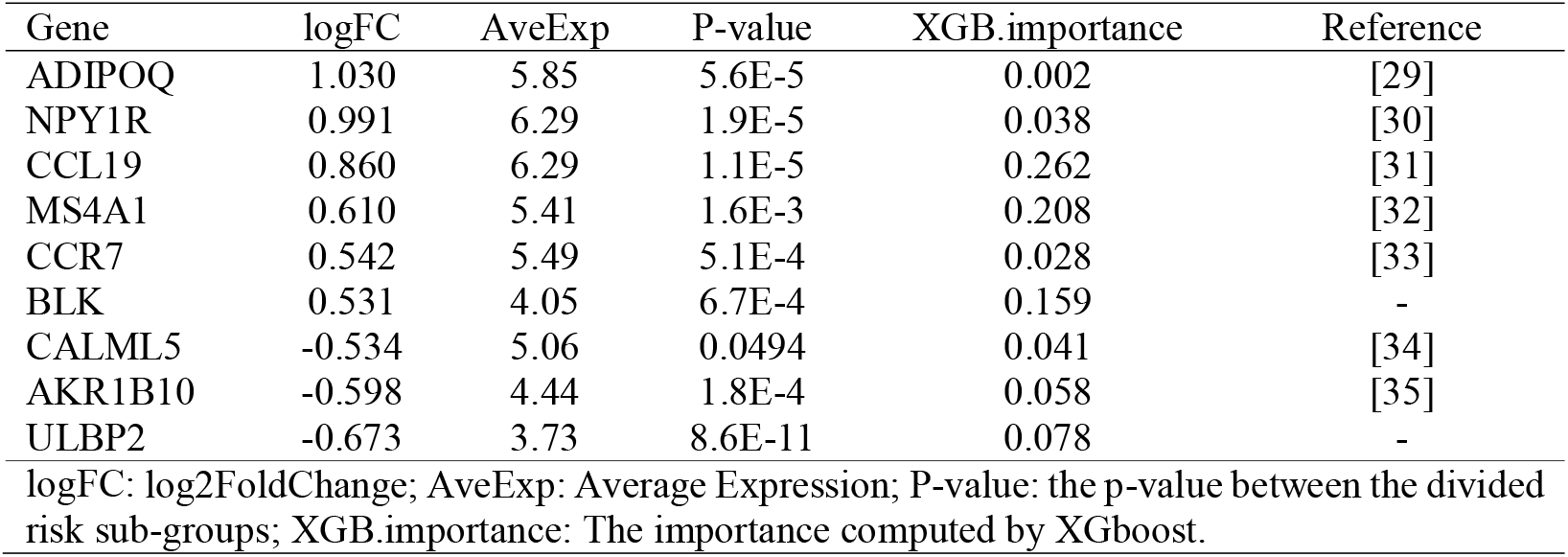
The identified prognostic markers in breast cancer

| Gene | logFC | AveExp | P-value | XGB.importance | Reference |
| --- | --- | --- | --- | --- | --- |
| ADIPOQ | 1.030 | 5.85 | 5.6E-5 | 0.002 | [29] |
| NPY1R | 0.991 | 6.29 | 1.9E-5 | 0.038 | [30] |
| CCL19 | 0.860 | 6.29 | 1.1E-5 | 0.262 | [31] |
| MS4A1 | 0.610 | 5.41 | 1.6E-3 | 0.208 | [32] |
| CCR7 | 0.542 | 5.49 | 5.1E-4 | 0.028 | [33] |
| BLK | 0.531 | 4.05 | 6.7E-4 | 0.159 | - |
| CALML5 | -0.534 | 5.06 | 0.0494 | 0.041 | [34] |
| AKR1B10 | -0.598 | 4.44 | 1.8E-4 | 0.058 | [35] |
| ULBP2 | -0.673 | 3.73 | 8.6E-11 | 0.078 | - |

logFC: $\log_2\text{FoldChange}$ ; AveExp: Average Expression; P-value: the p-value between the divided risk sub-groups; XGB.importance: The importance computed by XGboost.

## Conclusion and Discussion

Previously, many methods used individual types of genomic data to identify high-risk cancer patients from low-risk ones. Since individual omics type only offered a single view of cancers, the performances of these methods were limited. The multi-omics data analysis could bring more information about the cancer survival. In this study, we designed a deep learning framework DCAP to integrate the multi-omics data for cancer risk estimation. By comparing the prognosis prediction accuracy, the results obtained by DCAP outperformed the compared methods by > 6.5% C-index value on average. The ablation study showed that mRNA performs the best, the miRNA and methylation ranked the 2^nd^ and 3^rd^, and the copy number variation showed the least contribution. At last, in the case study on breast cancer, the independent tests of 3 GEO data proved that the constructed prediction model can separate the high-risk patients from the low-risk ones significantly (C-index>0.6, p-values<0.05). Based on the risk subgroups divided by DCAP, we identified nine prognostic markers highly associated with the breast cancer.

Though our method was indicated robust and reliable for predicting cancer outcomes, there are still many questions worth to be discussed below. Firstly, we found that the C-index values of different cancers obtained by using TCGA data fluctuated greatly. One possible reason is the ignoring of tumor purity and clinical factors that were already known to be important in TCGA [36]. Secondly, many censored samples in the data limited the accuracy of predicting cancer outcomes. For example, the censored rates are 60.7% and 64.5% for STAD and LUAD, respectively. The high censored rates decreased the performance of our method. Thirdly, one previous study [15] showed that the clinical data is helpful to improve the cancer prognosis prediction performance. This is a potential way to improve the model prediction performance.

In the future, we will consider the impact of heterogeneity caused by different clinical characteristics (including age and sex) on the prognostic risk of cancer patients. Additionally, we will optimize the neural networks by directly optimizing the risk loss function. At last, multi-modal medical data have been used for estimating cancer progress [15, 37]. We will further combine medical information such as slide images and clinical data for more accurate cancer prognosis estimation.

## List of abbreviations

AE-Cox: The Cox model using the compressed features by Autoencoder
BLCA: Bladder Urothelial Carcinoma
BRCA: Breast invasive carcinoma
CESC: Cervical squamous cell carcinoma and endocervical adenocarcinoma
CNV: Copy number variation
COAD: Colon adenocarcinoma
ConcatAE-Cox: The Cox model using the compressed features by ConcatAE
Cox-EN: Cox with elastic net
Cox-PH: Cox proportional hazard model
DCAP: A framework to integrate multi-omics data by Denoising Autoencoder for Accurate cancer prognosis prediction
DEG: Differentially expressed gene
DL-Cox: Deep neural network Cox method
ESCA: Esophageal carcinoma
GEO: Gene Expression Omnibus database
HC-Cox: A hierarchical clustering framework to extract multi-omics data for cancer prognosis prediction
HNSC: Head and Neck squamous cell carcinoma
LGG: Brain Lower Grade Glioma
LIHC: Liver hepatocellular carcinoma
LUAD: Lung adenocarcinoma
LUSC: Lung squamous cell carcinoma
MESO: Mesothelioma
PAAD: Pancreatic adenocarcinoma
PCA-Cox: Cox model used the reconstructed features by PCA
SARC: Sarcoma
SKCM: Skin Cutaneous Melanoma
STAD: Stomach adenocarcinoma
TCGA: The Cancer Genome Atlas

## Declarations

### Ethics approval and consent to participate

Not applicable

### Consent for publication

All the authors listed have approved the manuscript.

### Availability of data and materials

All the data analyzed during the current study are available in the TCGA dataset (https://tcga-data.nci.nih.gov/tcga/).

The method codes are available at https://github.com/Hua0113/DCAP

## Acknowledgements

We thank Mr. Ziwang Huang in Sun Yat-sen University for supporting our research with data collection.

## Authors’ contributions

CH and YY conceived the study. CH, ZX, and RJ performed the data analysis. CH, ZZ, ZH, and YY interpreted the results. CH, ZH, and YY wrote the manuscript. All authors read and approved the final manuscript.

## Supplementary Materials

**Table S1**. The contribution of each omics data for different cancer outcomes evaluation by using only one type of omics data or subtracting one type from the final model.

**Table S2**. The number of selected genes by DCAP-XGB in TCGA.

**Fig S1**. Convergence Analysis in DAE.

**Fig S2**. Parameter Sensitivity Study in DAE. (A) Learning rate. (B) Hidden node size in middle-hidden layer. (C) Batch size.

## References

[1] I. Dagogo-Jack, A.T. Shaw, Tumour heterogeneity and resistance to cancer therapies, Nat Rev Clin Oncol, 15 (2018) 81–94. https://doi.org/10.1038/nrclinonc.2017.166.

[2] K. Kourou, T.P. Exarchos, K.P. Exarchos, M.V. Karamouzis, D.I. Fotiadis, Machine learning applications in cancer prognosis and prediction, Comput Struct Biotechnol J, 13 (2015) 8–17. https://doi.org/10.1016/j.csbj.2014.11.005.

[3] L. Wang, Y. Li, J. Zhou, D. Zhu, J. Ye, Multi-task survival analysis, 2017 IEEE International Conference on Data Mining (ICDM), IEEE, 2017, pp. 485–494.

[4] C. Stirzaker, E. Zotenko, J.Z. Song, W. Qu, S.S. Nair, W.J. Locke, A. Stone, N.J. Armstong, M.D. Robinson, A. Dobrovic, K.A. Avery-Kiejda, K.M. Peters, J.D. French, S. Stein, D.J. Korbie, M. Trau, J.F. Forbes, R.J. Scott, M.A. Brown, G.D. Francis, S.J. Clark, Methylome sequencing in triple-negative breast cancer reveals distinct methylation clusters with prognostic value, Nat Commun, 6 (2015) 5899. https://doi.org/10.1038/ncomms6899.

[5] S. Volinia, C.M. Croce, Prognostic microRNA/mRNA signature from the integrated analysis of patients with invasive breast cancer, Proc Natl Acad Sci U S A, 110 (2013) 7413–7417. https://doi.org/10.1073/pnas.1304977110.

[6] Y. Wu, H. Chen, G. Jiang, Z. Mo, D. Ye, M. Wang, J. Qi, X. Lin, S.L. Zheng, N. Zhang, R. Na, Q. Ding, J. Xu, Y. Sun, Genome-wide Association Study (GWAS) of Germline Copy Number Variations (CNVs) Reveal Genetic Risks of Prostate Cancer in Chinese population, J Cancer, 9 (2018) 923–928. https://doi.org/10.7150/jca.22802.

[7] K. Tomczak, P. Czerwinska, M. Wiznerowicz, The Cancer Genome Atlas (TCGA): an immeasurable source of knowledge, Contemp Oncol (Pozn), 19 (2015) A68–77. https://doi.org/10.5114/wo.2014.47136.

[8] F. Rohart, B. Gautier, A. Singh, K.A. Le Cao, mixOmics: An R package for ‘omics feature selection and multiple data integration, PLoS Comput Biol, 13 (2017) e1005752. https://doi.org/10.1371/journal.pcbi.1005752.

[9] J. Mariette, N. Villa-Vialaneix, Unsupervised multiple kernel learning for heterogeneous data integration, Bioinformatics, 34 (2018) 1009–1015. https://doi.org/10.1093/bioinformatics/btx682.

[10] D. Kim, R. Li, A. Lucas, S.S. Verma, S.M. Dudek, M.D. Ritchie, Using knowledge-driven genomic interactions for multi-omics data analysis: metadimensional models for predicting clinical outcomes in ovarian carcinoma, J Am Med Inform Assoc, 24 (2017) 577–587. https://doi.org/10.1093/jamia/ocw165.

[11] A. Ahmad, H. Frohlich, Towards clinically more relevant dissection of patient heterogeneity via survival-based Bayesian clustering, Bioinformatics, 33 (2017) 3558–3566. https://doi.org/10.1093/bioinformatics/btx464.

[12] P. Coretto, A. Serra, R. Tagliaferri, Robust clustering of noisy high-dimensional gene expression data for patients subtyping, Bioinformatics, 34 (2018) 4064–4072. https://doi.org/10.1093/bioinformatics/bty502.

[13] D. Tong, Y. Tian, T. Zhou, Q. Ye, J. Li, K. Ding, J. Li, Improving prediction performance of colon cancer prognosis based on the integration of clinical and multi-omics data, BMC medical informatics, 20 (2020) 22. https://doi.org/10.1186/s12911-020-1043-1.

[14] Y. Li, F.X. Wu, A. Ngom, A review on machine learning principles for multi-view biological data integration, Brief Bioinform, 19 (2018) 325–340. https://doi.org/10.1093/bib/bbw113.

[15] A. Cheerla, O. Gevaert, Deep learning with multimodal representation for pancancer prognosis prediction, Bioinformatics, 35 (2019) i446–i454. https://doi.org/10.1093/bioinformatics/btz342.

[16] K. Chaudhary, O.B. Poirion, L. Lu, L.X. Garmire, Deep Learning-Based Multi-Omics Integration Robustly Predicts Survival in Liver Cancer, Clin Cancer Res, 24 (2018) 1248–1259. https://doi.org/10.1158/1078-0432.CCR-17-0853.

[17] T.-Y. Lee, K.-Y. Huang, C.-H. Chuang, C.-Y. Lee, T.-H. Chang, Incorporating deep learning and multi-omics autoencoding for analysis of lung adenocarcinoma prognostication, Computational Biology, 87 (2020) 107277. https://doi.org/10.1016/j.compbiolchem.2020.107277.

[18] L. Tong, J. Mitchel, K. Chatlin, M.D. Wang, Deep learning based feature-level integration of multi-omics data for breast cancer patients survival analysis, BMC medical informatics, 20 (2020) 1–12. https://doi.org/10.1186/s12911-020-01225-8.

[19] P. Vincent, H. Larochelle, Y. Bengio, P.-A. Manzagol, Extracting and composing robust features with denoising autoencoders, Proceedings of the 25th international conference on Machine learning, 2008, pp. 1096–1103.

[20] L. Wei, Z. Jin, S. Yang, Y. Xu, Y. Zhu, Y. Ji, TCGA-assembler 2: software pipeline for retrieval and processing of TCGA/CPTAC data, Bioinformatics, 34 (2018) 1615–1617. https://doi.org/10.1093/bioinformatics/btx812.

[21] N. Bokde, F. Martinez Alvarez, M.W. Beck, K. Kulat, A novel imputation methodology for time series based on pattern sequence forecasting, Pattern Recognit Lett, 116 (2018) 88–96. https://doi.org/10.1016/j.patrec.2018.09.020.

[22] M.E. Ritchie, B. Phipson, D. Wu, Y. Hu, C.W. Law, W. Shi, G.K. Smyth, limma powers differential expression analyses for RNA-sequencing and microarray studies, Nucleic Acids Res, 43 (2015) e47. https://doi.org/10.1093/nar/gkv007.

[23] N. Simon, J. Friedman, T. Hastie, R. Tibshirani, Regularization Paths for Cox’s Proportional Hazards Model via Coordinate Descent, J Stat Softw, 39 (2011) 1–13. https://doi.org/10.18637/jss.v039.i05.

[24] Chen T, Guestrin C. Xgboost: A scalable tree boosting system, Proceedings of the 22nd acm sigkdd international conference on knowledge discovery and data mining. (2016) 785–794. https://doi.org/10.1145/2939672.2939785.

[25] V. Van Belle, K. Pelckmans, S. Van Huffel, J.A. Suykens, Support vector methods for survival analysis: a comparison between ranking and regression approaches, Artif Intell Med, 53 (2011) 107–118. https://doi.org/10.1016/j.artmed.2011.06.006.

[26] N. Simon, J. Friedman, T. Hastie, R.J.J.o.s.s. Tibshirani, Regularization paths for Cox’s proportional hazards model via coordinate descent, Journal of statistical software, 39 (2011) 1. https://doi.org/10.18637/jss.v039.i05.

[27] H. Li, T. Lakshmikanth, C. Garofalo, M. Enge, C. Spinnler, A. Anichini, L. Szekely, K. Kärre, E. Carbone, G.J.C.c. Selivanova, Pharmacological activation of p53 triggers anticancer innate immune response through induction of ULBP2, Cell cycle, 10 (2011) 3346–3358. https://doi.org/10.4161/cc.10.19.17630.

[28] D.L. Petersen, J. Berthelsen, A. Willerslev-Olsen, S. Fredholm, S. Dabelsteen, C.M. Bonefeld, C. Geisler, A.J.T.B. Woetmann, A novel BLK-induced tumor model, Tumor Biology, 39 (2017) 1010428317714196. https://doi.org/10.1177/1010428317714196.

[29] S.J. Chung, G.P. Nagaraju, A. Nagalingam, N. Muniraj, P. Kuppusamy, A. Walker, J. Woo, B. Győrffy, E. Gabrielson, N.K.J.A. Saxena, ADIPOQ/adiponectin induces cytotoxic autophagy in breast cancer cells through STK11/LKB1-mediated activation of the AMPK-ULK1 axis, Autophagy, 13 (2017) 1386–1403. https://doi.org/10.1080/15548627.2017.1332565.

[30] L. Liu, Q. Xu, L. Cheng, C. Ma, L. Xiao, D. Xu, Y. Gao, J. Wang, H.J.O.l. Song, NPY1R is a novel peripheral blood marker predictive of metastasis and prognosis in breast cancer patients, Oncology letters, 9 (2015) 891–896. https://doi.org/10.3892/ol.2014.2721.

[31] H. Hwang, C. Shin, J. Park, E. Kang, B. Choi, J.-A. Han, Y. Do, S. Ryu, Y.-K. Cho, Human breast cancer-derived soluble factors facilitate CCL19-induced chemotaxis of human dendritic cells, Scientific reports, 6 (2016) 1–12. https://doi.org/10.1038/srep30207.

[32] A. Calabrò, T. Beissbarth, R. Kuner, M. Stojanov, A. Benner, M. Asslaber, F. Ploner, K. Zatloukal, H. Samonigg, A.J.B.c.r. Poustka, treatment, Effects of infiltrating lymphocytes and estrogen receptor on gene expression and prognosis in breast cancer, Breast cancer research and treatment, 116 (2009) 69–77. https://doi.org/10.1007/s10549-008-0105-3.

[33] N. Cabioglu, M.S. Yazici, B. Arun, K.R. Broglio, G.N. Hortobagyi, J.E. Price, A.J.C.C.R. Sahin, CCR7 and CXCR4 as novel biomarkers predicting axillary lymph node metastasis in T1 breast cancer, Clinical Cancer Research, 11 (2005) 5686–5693. https://doi.org/10.1158/1078-0432.CCR-05-0014

[34] M. Debald, F.A. Schildberg, A. Linke, K. Walgenbach, W. Kuhn, G. Hartmann, G.J.J.o.c.r. Walgenbach-Bruenagel, c. oncology, Specific expression of k63-linked ubiquitination of calmodulin-like protein 5 in breast cancer of premenopausal patients, Journal of cancer research and clinical oncology, 139 (2013) 2125–2132. https://doi.org/10.1007/s00432-013-1541-y.

[35] J. Ma, D.X. Luo, C. Huang, Y. Shen, Y. Bu, S. Markwell, J. Gao, J. Liu, X. Zu, Z.J.I.j.o.c. Cao, AKR1B10 overexpression in breast cancer: association with tumor size, lymph node metastasis and patient survival and its potential as a novel serum marker, International journal of cancer, 131 (2012) E862–E871. https://doi.org/10.1002/ijc.27618.

[36] D. Aran, M. Sirota, A.J. Butte, Systematic pan-cancer analysis of tumour purity, Nat Commun, 6 (2015) 8971. https://doi.org/10.1038/ncomms9971.

[37] P. Mobadersany, S. Yousefi, M. Amgad, D.A. Gutman, J.S. Barnholtz-Sloan, J.E. Velazquez Vega, D.J. Brat, L.A.D. Cooper, Predicting cancer outcomes from histology and genomics using convolutional networks, Proc Natl Acad Sci U S A, 115 (2018) E2970–E2979. https://doi.org/10.1073/pnas.1717139115.

